# FishResp: R Package and GUI Application for Analysis of Aquatic Respirometry Data

**DOI:** 10.1101/467845

**Authors:** Sergey Morozov, R. J. Scott McCairns, Juha Merilä

**Affiliations:** Ecological Genetics Research Unit, Organismal and Evolutionary Biology Programme, Faculty of Biological and Environmental Sciences, University of Helsinki, Finland.; ESE, Ecology and Ecosystem Health, INRA, Agrocampus Ouest, Rennes, France

**Keywords:** background respiration, free software, intermittent-flow respirometry, metabolic rate, oxygen uptake, three-spined stickleback, Trinidadian guppy

## Abstract

Intermittent-flow respirometry is widely used to measure oxygen uptake rates and subsequently estimate aerobic metabolic rates of aquatic animals. However, the lack of a standard quality-control software to detect technical problems represents a potential impediment to comparisons across studies. Here, we introduce ‘FishResp’, a versatile R package and its graphical implementation for quality-control and filtering of raw respirometry data. Our goal is to provide a straightforward, cross-platform and free software to help improve the quality and comparability of metabolic rate estimates for reducing methodological fragmentation in the field of aquatic respirometry. FishResp accepts data from various respirometry systems, allows users to detect potential mechanical problems which can occur during oxygen uptake measurements (e.g. chamber leaking, poor water circulation), and offers six options to correct raw data for microbial oxygen consumption. The software performs filtering of raw data based on user criteria, and produces accurate and unbiased estimates of absolute and mass-specific metabolic rates. Using data from three-spined sticklebacks (*Gasterosteus aculeatus*) and Trinidadian guppies (*Poecilia reticulata*), we demonstrate the virtues of FishResp, highlighting the importance of detecting mechanical problems and correcting measurements for background respiration.

**Lay summary:** FishResp is a user-friendly tool for calculating oxygen uptake of aquatic organisms. The aim of the software is to improve the quality of metabolic rate estimates based on a straightforward pipeline: background respiration correction, detection of mechanical problems, conduction of QC tests, and filtration based on user-defined criteria.

## 1 Introduction

Whole-body intermittent-flow respirometry (i.e. measuring the oxygen uptake of an organism) represents the most reliable and widespread technique to estimate aerobic metabolic rate of aquatic animals (Steffensen, 1989; Clark et al., 2013; Svendsen et al., 2016b). As the availability and precision of oxygen sensors has grown (Nelson, 2016), respirometry has become increasingly common in the field of evolutionary and conservation physiology (Chabot et al., 2016a; Metcalfe et al., 2016). Among other fundamental questions, it has been applied to the identification of ecological and environmental factors that cause metabolic rate variation (Ohlberger et al., 2007; Auer et al., 2015; Christensen et al., 2017) and for studying physiological plasticity and adaptation to novel environmental conditions, including adaptive potential to global ocean warming, ocean acidification, hypoxia and chemical contaminants (Kreiss et al., 2015; McDonnell and Chapman, 2015; Hancock and Place, 2016; Kunz et al., 2016; Norin et al., 2016; Jayasundara et al., 2017; Zhang et al., 2017). In addition, intermittent-flow respirometry has been recently used in studies devoted to both applied and fundamental problems in aquaculture and fisheries (Hessenauer et al., 2015; Siikavuopio and James, 2015; Gerile and Pirhonen, 2017; Svendsen et al., 2018).

Despite the growing use of respirometry, quality-control of raw data and extraction of key parameters from it, remains the responsibility of individual researchers (Rodgers et al., 2016). The lack of standard quality-control software with the ability to detect and correct for technical problems in the raw data may compromise the comparability of results from different studies. Indeed, the literature contains many contradictory reports of intraspecific metabolic differences that are difficult to ascribe to only biological causes. For example, whilst inter-population comparisons of mean standard metabolic rate (SMR) estimates of Trinidadian guppy (*Poecilia reticulata*) have reported difference up to 1.7-fold between populations (Handelsman et al., 2013; Auer et al., 2018), comparisons between independent studies conducted under similar temperature conditions show 5-fold differences (from 100 to 550 mg O_2_/kg/h) (Svendsen et al., 2013; Ejbye-Ernst et al., 2016; Killen et al., 2016). Such a wide range of SMR estimates within a species are unlikely to be explanaible by biological factors, but more likely, reflect unstandardized measurement conditions and data post-processing procedures.

Two critical factors that influence the quality and reliability of respirometry data are (i) mechanical problems with the system itself, and (ii) background respiration. As to the former, both the absence of or damage to a mixing device leading to poor water circulation in a system, and a large ratio of respirometer to fish volume, can reduce the resolution of raw data (Rodgers et al., 2016; Svendsen et al., 2016a). Similarly, study animals blocking the water supply to an oxygen sensor can cause dramatic drops in dissolved oxygen measurements, whereas leaking of a measurement chamber, which can be particularly pronounced under high water flow velocity in Blazka-type swimming tunnels (Blazka et al., 1960), has potential to lead to underestimation of metabolic rates (Svendsen et al., 2016b). Mechanical problems are relatively easy to detect via linear analysis of raw data, and may indicate the need to repair faulty equipment. However, detecting and correcting for background respiration (i.e. oxygen consumption of microorganisms in the water or on inner surfaces of equipment) can be more challenging (Dalla Via, 1983; Clark et al., 2013; Rodgers et al., 2016). Although several approaches exist, many studies fail to mention how background respiration was dealt with, or whether background respiration was totally ignored in data processing (see Svendsen et al., 2016b). The potential problem with background respiration is especially relevant when metabolic rate is measured at high temperatures – a condition that favors microbial growth – and in small chambers, where the relative contribution of microbes to total rate of oxygen consumption is high (Rodgers et al., 2016). Both are prevalent when working with small tropical fish, as in the aforementioned guppy example, and can lead to substantial overestimates of oxygen consumption rate. In addition, background respiration may represent an increasingly large proportion of the total oxygen consumption under hypoxic conditions.

Here, we introduce ‘FishResp’, software for running quality control tests, filtering raw data, and calculating metabolic estimates of high accuracy. We demonstrate the utility of FishResp for detecting mechanical problems and correcting for background microbial oxygen consumption using raw data obtained from the three-spined stickleback (*Gasterosteus aculeatus*) and the Trinidadian guppy (*Poecilia reticulata*). These examples demonstrate how FishResp significantly improves the quality of final estimates, setting what we hope will become standard practice for processing raw respirometry data.

## 2 Materials and Methods

### 2.1 Software Description

FishResp is an R package (R Development Core Team, 2015) for analyzing raw data from intermittent-flow respirometry systems. Its current version (1.0.2 consists of 15 functions (listed in Table 1) and utilizes five dependencies (existing R packages) which will be automatically installed once FishResp is executed (listed in Table 2). The order of application of each function in a typical analysis is described in Figure 1, encompassing all steps from data input to parameter estimation. Alternatively, a graphical user interface (GUI) of FishResp is also available, written in JavaFX. To integrate Java with R, the ‘rJava’ package (Urbanek, 2018) and its extension ‘JRI’ are used. The graphical interface consists of five tabs with functional blocks on the left side (functions with parameters), and information, graphical or extensions modules on the right side of the application (Fig. 2). The ‘Extensions Module’ has been created to integrate FishResp with other respirometry tools written in the R language, such as modified functions for unit conversion from the R packages ‘respirometry’ and ‘rMR’ (Birk, 2018; Moulton, 2018). Different colors of tabs and buttons are used as indicators of progress: unavailable (grey); plotting optional (blue); action required (yellow); successful execution (green). The log of complete actions can be saved using an icon in the upper-right corner of the graphical interface. Both versions of FishResp are free and open-source software, compatible with Windows, MacOS and Linux.

**Figure 1.**
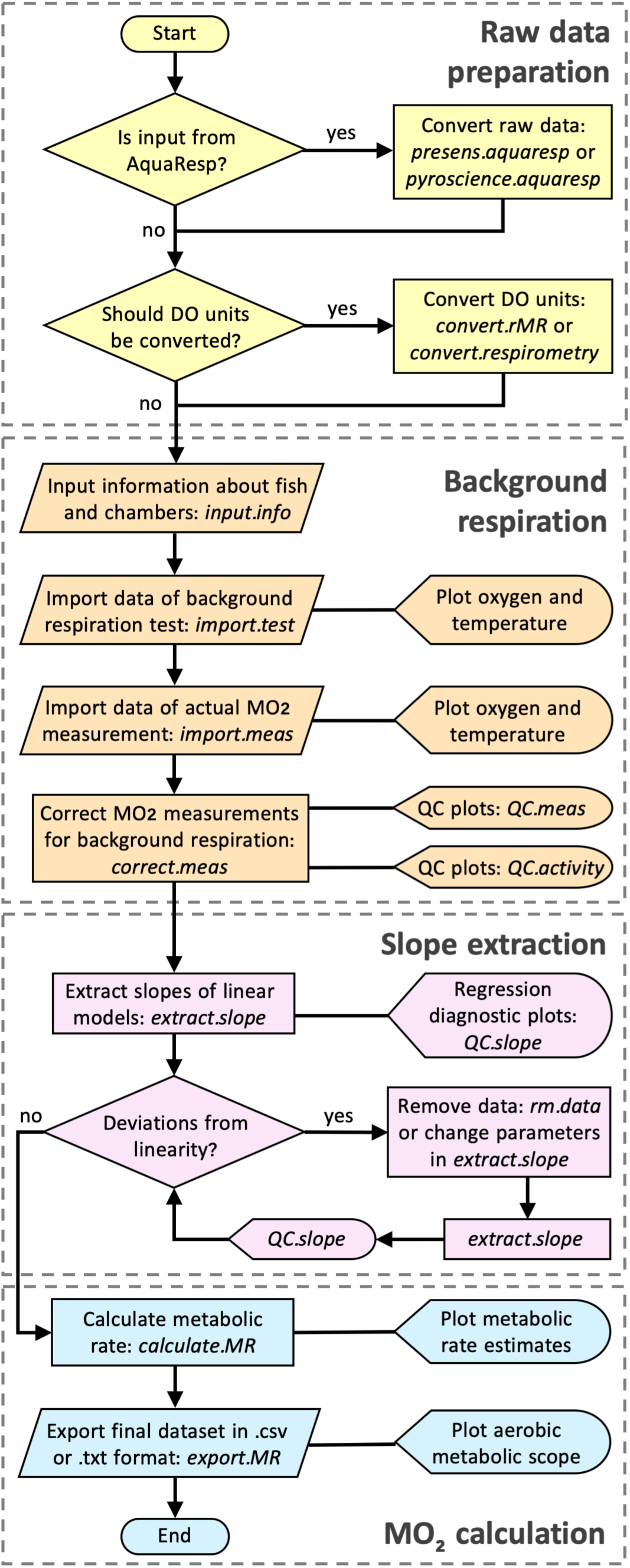
Flowchart illustrating a data processing pipeline for conducting analysis in the R package ‘FishResp’. The names of R functions are italicized. The functions are grouped into four categories: raw data preparation (optional), input data and background respiration correction, slope extraction based on parameter filtration, and calculation of metabolic rate estimates.

**Figure 2.**
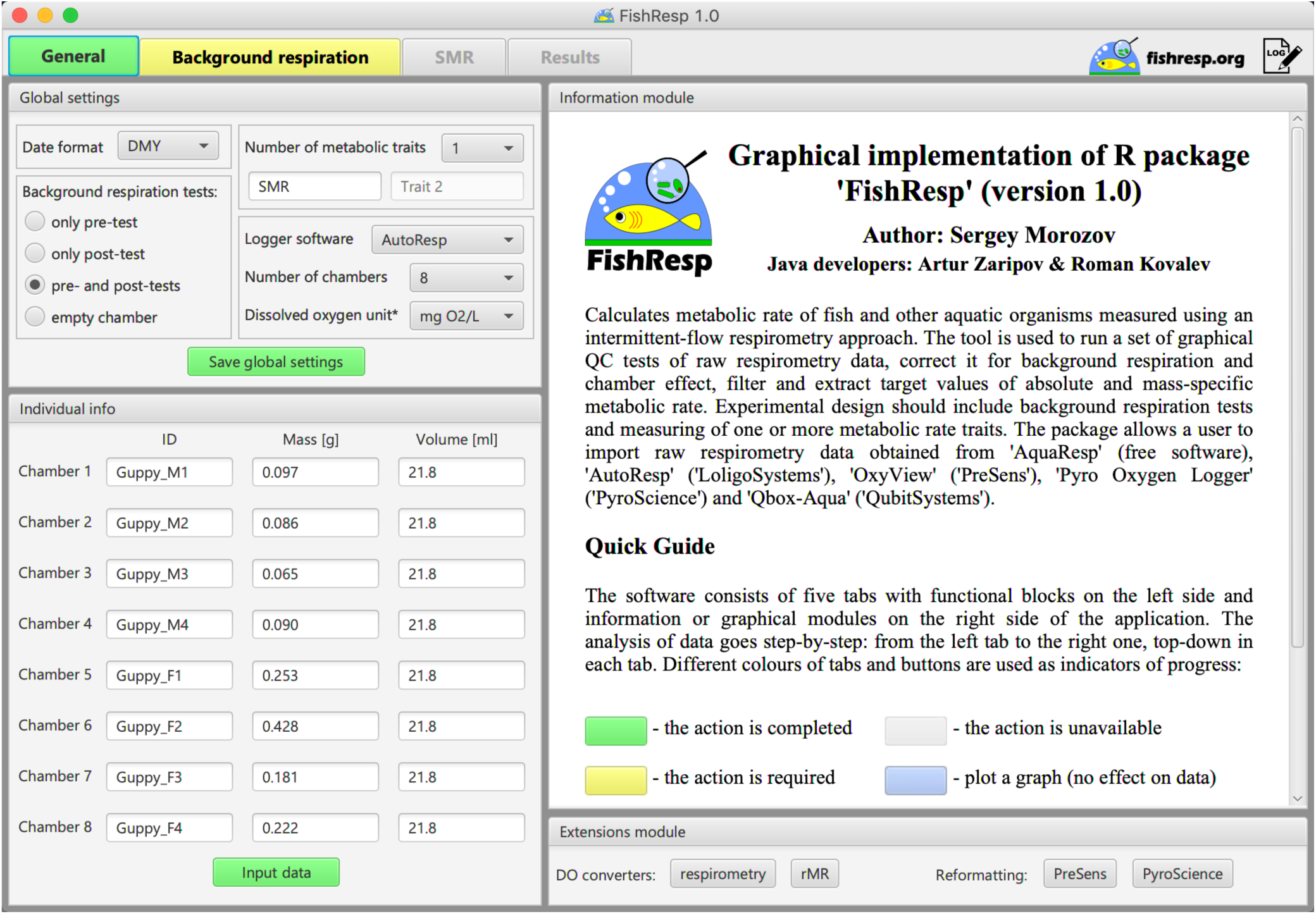
GUI implementation of ‘FishResp’: general settings and the information about guppies used in Case Study 2

**Table 1.** Overview of all functions in the R package. The functions are listed in order of a typical FishResp analysis (see also Fig.1), excluding optional functions *convert.respirometry*, *convert.rMR*, *presens.aquaresp*, *pyroscience.aquaresp*.

| Function | Description |
| --- | --- |
| *input.info* | Input information about experimental organisms and chambers |
| *import.test* | Import raw data for a background respiration test |
| *import.meas* | Import raw data for metabolic rate measurements |
| *correct.meas* | Correction of measurements for background respiration |
| *QC.meas* | Graphical quality control of temperature and O<sub>2</sub> measurements before and after correction for bacterial respiration |
| *QC.activity* | Graphical quality control of measurement windows |
| *extract.slope* | Extraction of target measurements |
| *QC.slope* | Graphical quality control of extracted measurements |
| *rm.data* | Remove raw data manually for a specific measurement phase |
| *calculate.MR* | Calculation and visualisation of background respiration, absolute and mass-specific metabolic rates |
| *export.MR* | Export results as .csv and .txt files (optionally, calculate aerobic metabolic scope) |
| *convert.respirometry*<br>& *convert.rMR* | Convert dissolved oxygen to partial pressure units and *vice versa* in raw data obtained in multichannel respirometry systems (see the R packages ‘respirometry’ and ‘rMR’) |
| *presens.aquaresp* &<br>*pyroscience.aquaresp* | Convert raw data from OxyView (PreSens) or Pyro Oxygen Logger (PyroScience) along with a summary file from AquaResp (free software) to the FishResp format |

**Table 2.** Overview of package dependencies and its functions in FishResp

| **R package** | **Function** | **Reference** |
| --- | --- | --- |
| *chron* | time analysis | (James and Hornik, 2018) |
| *lattice* | graphics | (Sarkar, 2008) |
| *mclust* | SMR determination | (Scrucca et al., 2016) |
| *respirometry* | DO converter | (Birk, 2018) |
| *rMR* | DO converter | (Moulton, 2018) |
| *rJava* | R to Java Interface | (Urbanek, 2018) |

The software allows a user to analyze data from up to eight respirometry chambers concurrently. Raw data can be imported into the R package/GUI application directly from ‘AutoResp’ (Loligo Systems, Denmark) and ‘Q-box Aqua (Qubit Systems, Canada); and via ‘AquaResp’ (University of Copenhagen, Denmark) with ‘OxyView’ (PreSens GmbH, Germany) and ‘Pyro Oxygen Logger’ (Pyro Science GmbH, Germany) output. The output from other logger software and DIY setups can be manually adjusted to ‘FishResp’ import format (see ‘Details’ in the documentation of the R function ‘import.test’ or ‘import.meas’). The software accepts several units of O_2_ concentration, while partial pressure units (e.g. % sat, mmHg, or kPa) can be converted to O_2_ concentration using the R functions ‘convert.respirometry’ and ‘convert.rMR’ or in the ‘Extensions Module’ of the GUI application (for details see the documentation of those functions). Quality-control tests and data filtering are performed with the aid of graphical visualizations, and parameters from simple linear models: the slope of the linear regression of O_2_ concentration over time, and the coefficient of determination (*r^2^*). Slopes are used to calculate metabolic rate, and to correct for background respiration.

### 2.2 Experimental Design for Correction of Background Respiration

The number of traits calculated per one session is not limited in the R package, while GUI implementation of FishResp can be used only for measuring one or two metabolic rates per session, as shown in Figure 3. In order to subtract background respiration from total oxygen consumption, additional tests must be conducted without an animal in the chambers before and/or after metabolic rate measurements (pre- and post-tests, respectively). A full experiment processed with the GUI application can consist of four different steps (Fig. 3A), but this can be reduced to one metabolic measurement with either one or two background respiration tests (Fig. 3B). Both full and reduced designs are described to illustrate the use of the R package and its GUI version.

**Figure 3.**
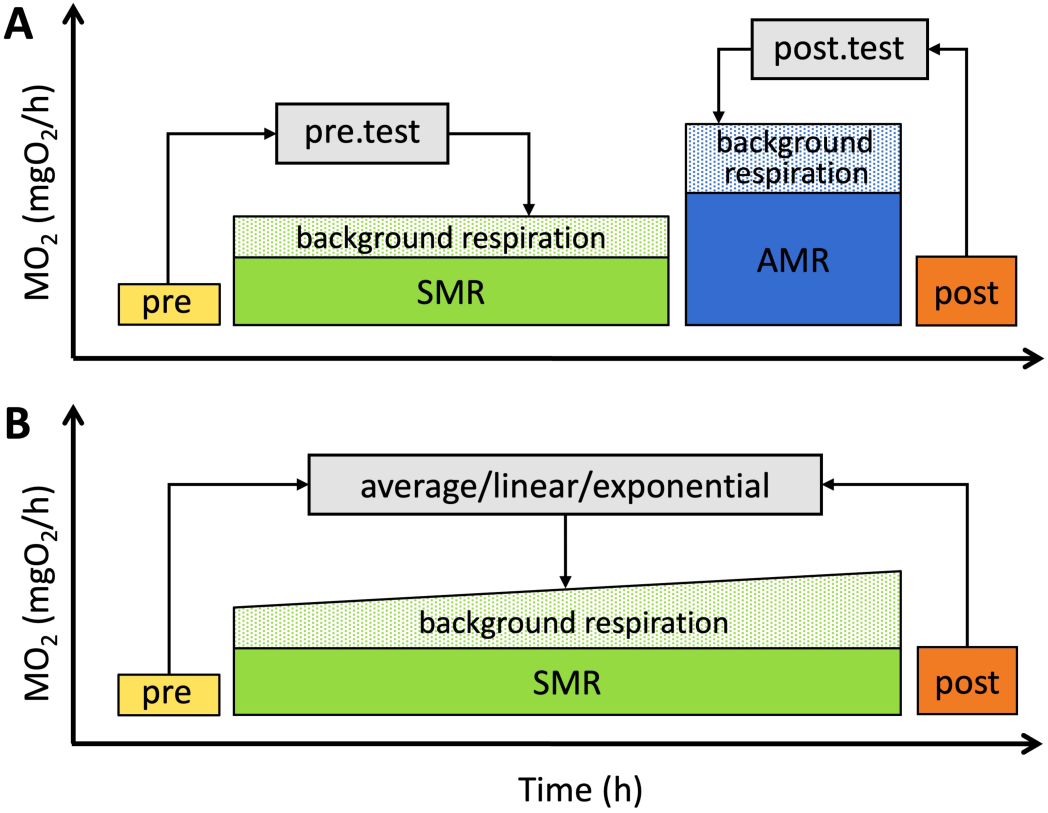
Schematic representation of full (A) and reduced (B) experimental designs. SMR and AMR are conditional terms and can be substituted with other traits. Pre and post blocks represent tests of background oxygen consumption before and after experiments.

Six options are available for the correction of background respiration, each based on subtracting the slope of background respiration over time from the slope of animal oxygen consumption over time. Methods ‘pre.test’ and ‘post.test’ are used to remove background respiration before or after metabolic rate measurements, respectively. When a single metabolic trait has been measured with both pre- and post-tests (Fig. 3B), ‘average’, ‘linear’ and ‘exponential’ methods can be used to subtract the averaged slope of the tests, or the progressively changing slope over time between those tests. Alternatively, an empty chamber running in parallel with actual metabolic rate measurements can be used for the correction. Note that the ‘parallel’ method does not require running pre- and/or post-tests. To choose an optimal method of correction, monitoring of background respiration over a long measurement period is recommended (Rodgers et al., 2016; Svendsen et al., 2016b).

### 2.3 Metabolic Rate Calculation

Several methods of slope extraction are available to estimate metabolic rate: all slopes, slopes with minimal or maximal values, or those of a given quantile within the frequency distribution of observed slopes. In addition, four methods for determination of SMR were integrated into FishResp: quantiles, MLND, low10, and low10% (see Chabot et al., 2016b). Likewise, users can define a threshold *r^2^* value (default value of 0.95), the parameter used to detect mechanical problems in a respirometry system (e.g. chamber leaking and poor water flow) or spontaneous animal activity during SMR measurements (but see Clark et al., 2013; Norin and Clark, 2017; Zhang and Gilbert, 2017).

Calculation of absolute (absMO_2_; ‘cO_2_ unit’/h) and mass-specific (massMO_2_; ‘cO_2_ unit’/kg/h) metabolic rates are based on the following equations (Claireaux and Lagardère, 1999):

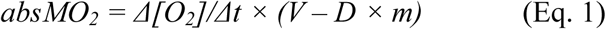

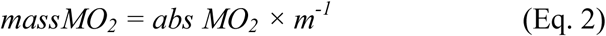

where *Δ[O_2_]/Δt* is slope in ‘cO_2_ unit’/L/h, *m* is mass in kg, *V* the chamber volume in L, and *D* is the density of an animal body (by default = 1000 kg/m^3^, can be modified; but see Svendsen et al. (2016b))

### 2.4 Fish and Respirometry Systems Used for Case Studies

Use of FishResp is demonstrated in two case studies. Case Study 1 shows how the full scheme can be run using the R package, and provides an example of how mechanical problems can be detected. Raw respirometry data come from four three-spined sticklebacks (average mass = 1.96, S.D. = 0.19 g), measured in a 4-channel respirometry system of 250 mL Blazka-type swim tunnels (DAQ-PAC-F4 package, AutoResp user’s manual; Loligo Systems, Denmark), together with pre- and post-tests.

Standard metabolic rate (SMR) was measured overnight (22:00-06:00) after four hours of acclimation in the absence of light and other external stimuli; active metabolic rate (AMR) was measured during the day at 80% of individual critical swimming speed, which represents a proxy of maximum aerobic activity during prolonged swimming performance, but not absolute maxMO_2_ (for definitions, see Brett, 1964; Fry, 1971). We used an intermittent-flow procedure consisting of flush, wait and measurement phases (420/180/1200 s for SMR and pre-test, and 420/180/600 s for AMR and post-test). The experiment was conducted in fresh water at 16.5°C with constant velocity (5 cm/s) generated by a propeller inside a swim tunnel. The level of dissolved oxygen during an open phase was 8.8 mg/L (90% air saturation), and it never dropped below 5.95 mg/L (61% air saturation) during the measurement phase.

Case Study 2, using data from four male and four female guppies (average mass_male_ = 0.084, S.D. = 0.014 g and average mass_female_ = 0.271, S.D. = 0.109 g), provides an example of excessively high background respiration and is presented to demonstrate both the GUI and the importance of background respiration correction. Measures of SMR were taken overnight after four hours of acclimation using eight 21.8 mL static chambers connected to an 8-channel respirometry system (DAQ-PAC-WF8 package, AutoResp user’s manual; Loligo Systems, Denmark) with 300 s flush, 180 s wait and 720 s measurement periods. Fresh water was constantly mixed by a pump during the measurement phase (T = 24.8°C; mean cO_2_ = 7.1, S.D. = 0.5 mg/L). Background respiration in the chambers was measured before and after the experiment (Fig. 3B). The calibration of the oxygen sensors for case studies was performed following the 2-point calibration procedure described in the AutoResp user’s manual (Loligo Systems, Denmark). Raw data for both Case Study 1 and Case Study 2 can be found in the package folders “FishResp/extdata/stickleback” and “FishResp/extdata/guppy”, respectively.

Experiments were carried out in accordance with the recommendations and approval of the National Animal Experiment Board of Finland (permit no: ESAVI/6097/04.10.07/2013) and the Stockholm Ethical Board (permit no: N173/13, 223/15 and N8/17).

## 3 Results and Discussion

### 3.1 Case Study 1: Calculation of SMR, AMR and MS of Sticklebacks

#### 3.1.1 Import and Correction of Raw Data

Following the general outline of a typical ‘FishResp’ analysis (Fig. 1), the command library(FishResp) is used to load the package. The function input.info() must be applied first to specify dissolved oxygen measurement units, and to provide information about individual ID, fish mass (g), and chamber volumes (mL) used in subsequent analyses (see Table 1):

~~~
info <- input.info(DO.unit = “mg/L”,
             ID = c(“Stickleback_1”, “Stickleback_2”, “Stickleback_3”, “Stickleback_4”),
             Mass = c(1.86, 1.92, 2.23, 1.80),
             Volume = c(250, 250, 250, 250))
~~~

To load files containing raw data of background respiration measurements into R (‘pre-test.txt’ and ‘post-test.txt’), the function import.test() should be used - note that if data files are not located in the same directory as the R session, file names should be specified with their full paths (see code in File S1). The path to the file, the type of oxygen logger software, and the number of chambers used should also be included as arguments. Graphs of oxygen and temperature over time are plotted automatically to check temperature fluctuations and water supply to the sensor during background respiration tests:

~~~
pre <- import.test(pre.path, info.data = info, logger = “AutoResp”, n.chamber = 4)
post <- import.test(post.path, info.data = info, logger = “AutoResp”, n.chamber = 4)
~~~

The next step is to import raw data of SMR and AMR measurements into R using the import.meas() function. If the date format is not ‘DMY’, specify it in this function. Two additional parameters (start.measure and stop.measure, respectively) corresponding to the time interval of measurements not exceeding 24 hours can help remove non-target data (e.g. acclimation period after handling stress):

~~~
SMR.raw <- import.meas(SMR.path, info.data = info, logger = “AutoResp”, n.chamber = 4,
                 start.measure = “22:00:00”, stop.measure = “06:00:00”)
AMR.raw <- import.meas(AMR.path, info.data = info, logger = “AutoResp”, n.chamber = 4)
~~~

As a full design was chosen for this experiment (Fig. 3A), ‘pre’ and ‘post’ methods are used to correct SMR and AMR for background respiration, respectively, where slopes of linear models (level of dissolved oxygen over time) for background respiration tests are automatically subtracted from slopes of linear models for actual measurements using the function correct.meas():

~~~
SMR.clean <- correct.meas(info.data = info, pre.data = pre,
                 meas.data = SMR.raw, method = “pre.test”)
AMR.clean <- correct.meas(info.data = info, post.data = post,
                 meas.data = AMR.raw, method = “post.test”)
~~~

Oxygen data before and after background respiration correction can be visualized using the function QC.meas():

~~~
QC.meas(SMR.clean, “Total.O_2_.phases”)
QC.meas(SMR.clean, “Corrected.O_2_.phases”)
QC.meas(SMR.clean, “Total.O_2_.chambers”)
QC.meas(SMR.clean, “Corrected.O_2_.chambers”)
~~~

Plotting raw data plays an important role in assessing its quality, providing preliminary information about the rate of background respiration; corrected O_2_ plots allow the user to evaluate their choice of correction strategy. In this example, levels of dissolved oxygen and temperature were stable over the duration of the experiment, indicating that both pre- and post-tests can be used to remove background respiration.

#### 3.1.2 Extraction and Visualization of Results: Identifying and Removing Artefacts due to Animal Disturbance or Swimming Inactivity

The next step is to select slopes of dissolved oxygen concentration over time to calculate metabolic rate. This is particularly relevant when several consecutive measurement cycles are taken. In this example, SMR measurements were collected every 30 min over an 8 h period, but only three slopes have been extracted to determine SMR. In contrast, as only three AMR measurements are available for each individual, all three slopes are used for calculation of AMR. For both traits, only slopes with *r^2^* > 0.95 are extracted.

~~~
SMR.slope <- extract.slope(SMR.clean, method = “min”, n.slope = 3, r^2^ = 0.95)
AMR.slope <- extract.slope(AMR.clean, method = “all”, r^2^ = 0.95)
~~~

It can be helpful to plot each slope separately: the function QC.slope() visualizes how a linear model fits data corrected for background respiration. Note, that an exponential decrease of oxygen content over long measurement periods might be caused by chamber leaking or poor water circulation in a respirometry system even with high *r^2^* value (Svendsen et al., 2016b; Norin and Clark, 2017). Would this be the case, those data should be omitted from further analyses.

~~~
QC.slope(SMR.slope, SMR.clean, chamber = “CH1”, current = 1200, alter = 600)
QC.slope(AMR.slope, AMR.clean, chamber = “CH4”, current = 600, alter = 300)
~~~

In this example, linear models perfectly capture the trend in SMR and most AMR data (Fig. 4, M1). However, data from the two final AMR measurement phases in ‘Chamber 4’ show decreases in oxygen uptake rate with time (Fig. 4, M2 & M3). During experiments (including Morozov et al., 2018), we noticed that some individuals could not constantly maintain 80% of U_crit_ in swimming tunnels during the final minutes of AMR measurements, but leaned against the honeycomb stopper housing the oxygen sensor. Occasionally, fish were able to block the oxygen supply to the sensor leading to dramatic decrease of oxygen level in raw data (Fig. 4, M2). The slopes for M2 look similar because blocking of the sensor was short and in the middle stage of measurement phase. The other common problem, when an experimental organism stopped swimming and rate of oxygen uptake dropped, is illustrated in panel M3 (Fig. 4, M3): the fish was not able to swim after first five minutes of the measurement phase, reflected in the upward inflection of the QC plot. In fact, here we detected both these mechanical problems based on graphical QC tests, but not based on r^2^ < 0.95. Thus, it is important to combine computational methods with visualization of intermittent results of the analysis. To avoid under- and overestimating AMR, and to follow a uniform approach for extracting slopes for this trait, we decreased the time interval used for slope calculation with the function extract.slope(). The other important feature of the function QC.slope(), which can represent a useful tool for detecting deviations from linearity and in determination of measurement window length, is regression diagnostic plots for current and alternative length of a measurement window (Fig. 5). In this example, reducing the measurement window to 5 min (300 s) helps improve linear fit and allow us to avoid artefacts due to swimming inactivity (Fig. 4; red lines, Fig. 5; red graphs):

~~~
QC.slope(AMR.slope, AMR.clean, chamber = “CH4”, current = 600, alter = 300, residuals = TRUE)
AMR.slope <- extract.slope(AMR.clean, method = “all”, r^2^ = 0.95, length = 300)
~~~

**Figure 4.**
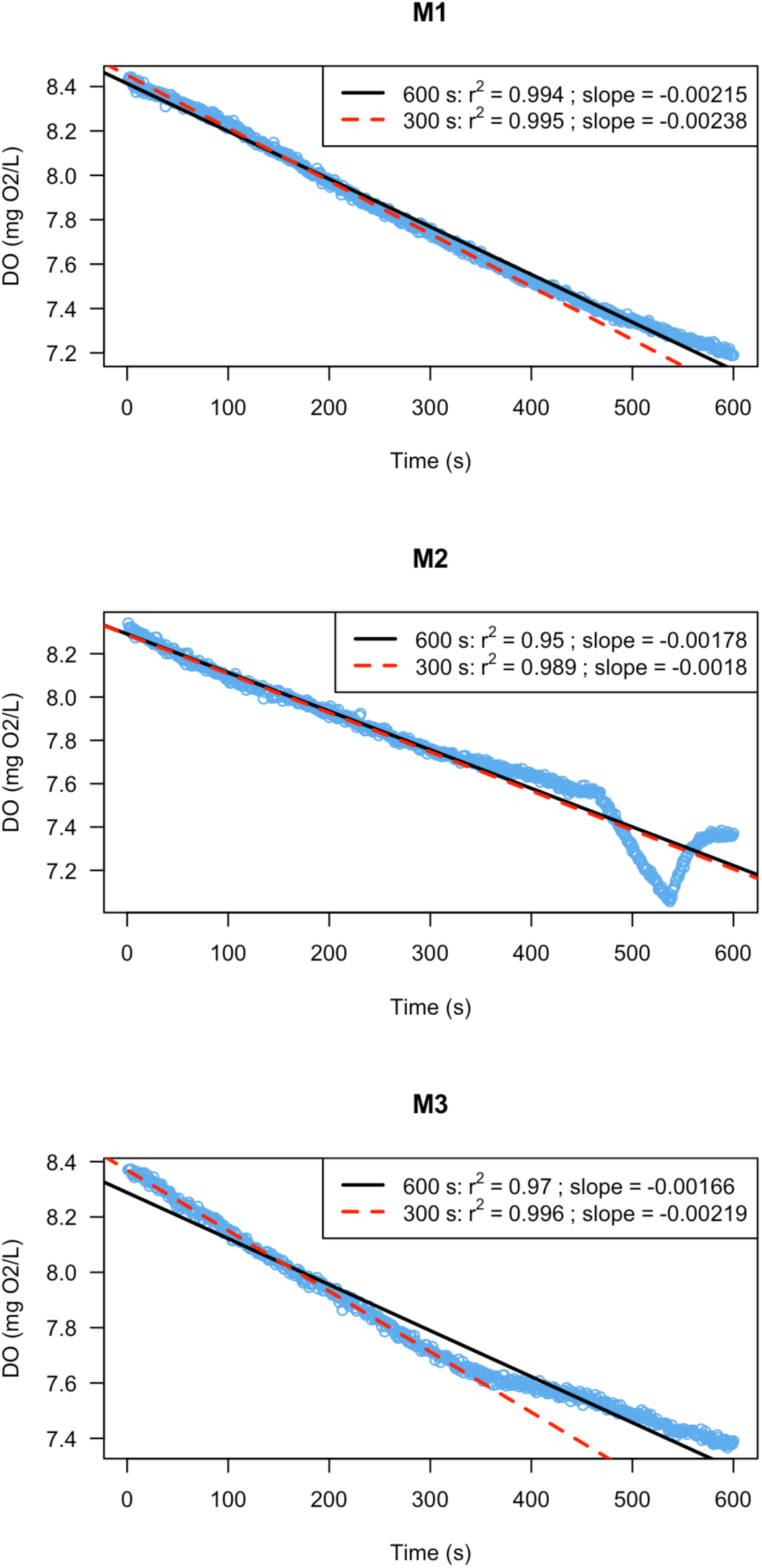
Slopes for AMR in three-spined stickleback. The level of oxygen content in a respirometry chamber was measured every second over 10 minutes during measurement phases - M. Black lines denote a linear model over the whole measurement period; in red measurements are restricted to the first five minutes. M1, M2 and M3 are the discrete measurement cycles.

**Figure 5.**
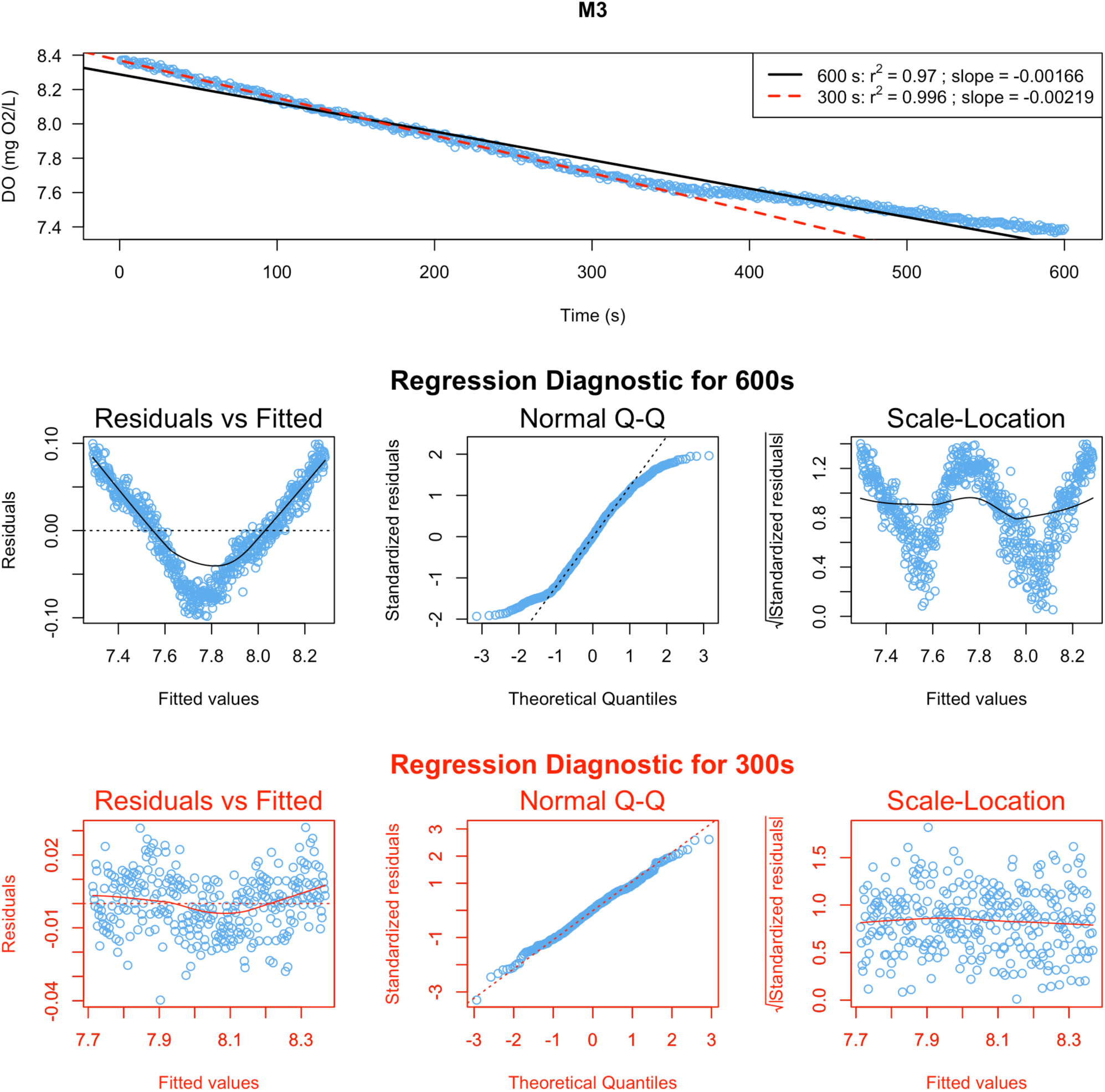
Regression diagnostics for the measurement phase ‘M3’ of Chamber 4. Raw data with fitted linear models for current length of measurement window (600s, black color) and the alternative one (300s, red color) displayed on the top graph. Regression diagnostic plots for current and alternative measurement windows are located in the middle section and the bottom section of the figure, respectively. More information on diagnostic plots can be found in the documentation of the base R function plot.lm().

It should be noted that a call to extract.slope() uses r^2^=0.95 as a default linearity filter. Svendsen et al. (2016b) have suggested that this should be viewed as the minimum criterion when estimating O_2_ consumption from the slope of a linear model. This value can be manually changed if users wish to use a more robust value. Under some circumstances users may wish to relax this parameter; however, we would urge careful reflection before doing so, and that all parameters used be reported.

The final step is to calculate and plot metabolic rate estimates, together with the percentage rate of background respiration using the function calculate.MR().

~~~
SMR <- calculate.MR(SMR.slope, density = 1000)
AMR <- calculate.MR(AMR.slope, density = 1000)
~~~

Results can be exported as .txt or .csv files. Optionally, when two traits are measured, they can be merged into one dataset containing absolute, mass-specific and factorial metabolic scopes.

~~~
results <- export.MR(SMR, AMR, “results.csv”, simplify = TRUE, MS = TRUE)
~~~

Note that the rate of background respiration for both SMR and AMR is not more than 13.8 % and 28.7 % respectively. However, the situation can change dramatically when the experiment is conducted at high water temperature and/or in small chambers, as demonstrated by Case Study 2.

### 3.2 Case Study 2: The Effect of Background Respiration on SMR of Guppies

#### 3.2.1 Running the Analysis Using GUI Application

The GUI version is created for users who prefer to interact with graphical elements in a user-friendly interface instead of working with a command line. Here, we demonstrate the use of the application for estimating background respiration, and calculating SMR (see File S2 to run this example in R). After launching the program, a user should set parameters in the ‘General Settings’ module: we choose “pre- and post-tests” as background respiration tests, ‘AutoResp’ for the type of logger software, and eight chambers in the respirometry system. As we measured only one trait (SMR), the field for ‘Trait 2’ should be empty. Click the ‘Save global settings’ button, and fill in the table with information about animals and chambers (Fig. 2). Next, move to the ‘Background respiration’ tab and import data for pre- and post-tests. Note that conducting visual QC tests of raw data is critical: if the temperature is unstable, or oxygen data show loss of linearity, further calculations might be meaningless. Navigation buttons of the graphical module can be used for zooming and switching plots for different chambers. Finally, open the ‘Standard Metabolic Rate’ tab, import raw data for SMR (time interval: 22:00:00 – 06:00:00), and correct them for background respiration. As both pre- and post-tests were used, all methods excluding ‘parallel’ are available (Fig. 3B). In this example, the method ‘exponential’ is used because background respiration was on average 17% lower during the pre-test than during the post-test in all chambers (Wilcoxon signed-rank test, *p* = 0.014), wherein exponential microbial growth is expected (Dalla Via, 1983; but see Rodgers et al., 2016).

#### 3.2.2 Comparison of SMR Before and After Correction

In the module ‘Visual QC tests’, a user can compare non-corrected versus corrected data, and check for indications of animal activity over the measurement period (GUI implementation of the package functions, QC.meas and QC.activity). ‘Activity’ and ‘Comparison’ options calculate mass-specific metabolic rate automatically, disabling any filters for slope estimation (but see Wilson, 2018). The ‘Activity’ option can be helpful for defining the time of corrected SMR, while ‘Comparison’ is used for graphical comparison of results with and without correction for background respiration (Fig. 6). Figure 6 shows that mass-specific SMR after the subtraction of microbial oxygen consumption is markedly lower than before correction.

**Figure 6.**
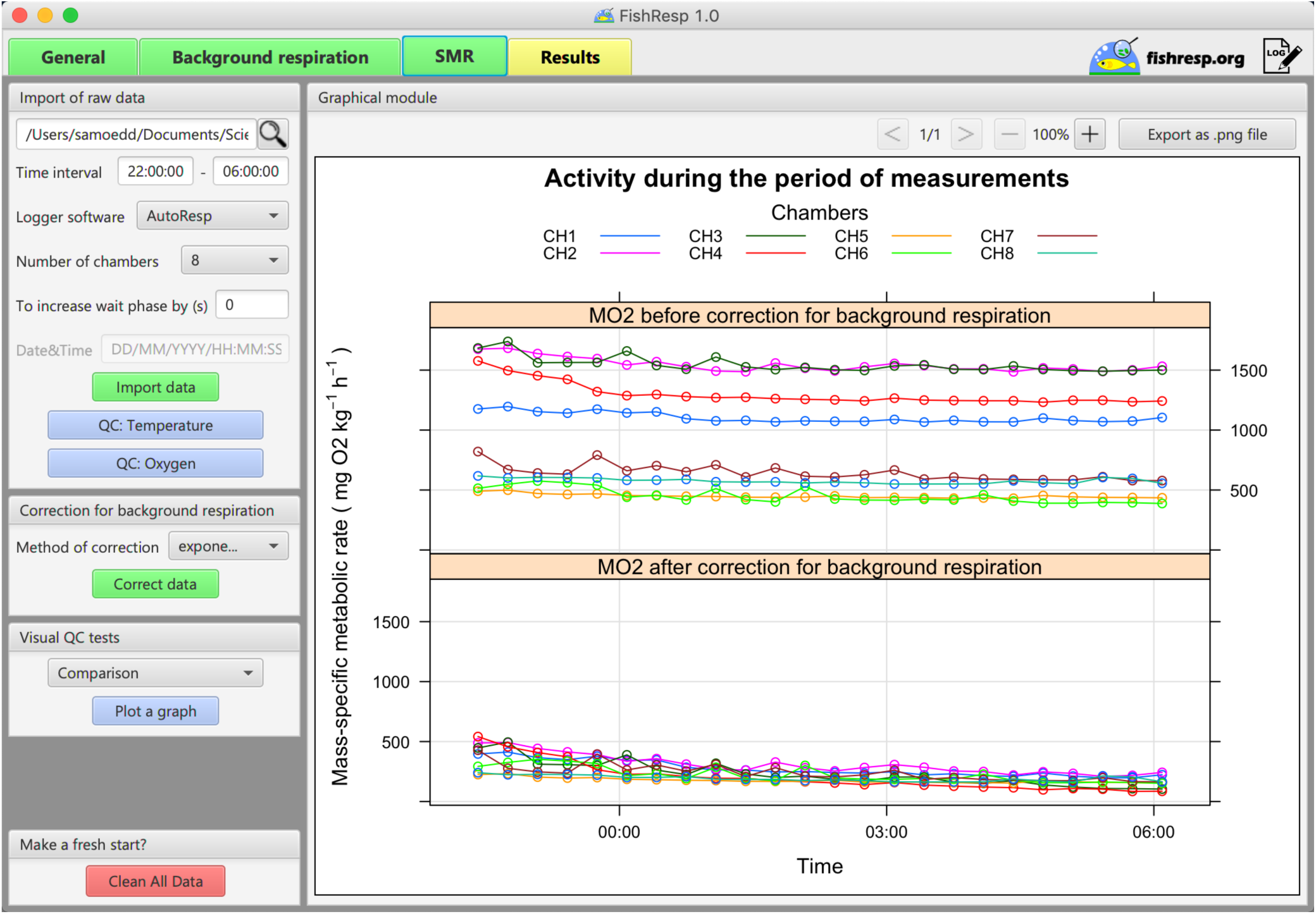
GUI implementation of ‘FishResp’: graphical module shows the difference in mass-specific SMR of guppies before (upper panel) and after correction (lower panel) for background respiration.

To calculate SMR, together with the percentage of background respiration, open the ‘Results’ tab. The method ‘calcSMR: quant’ (Chabot et al., 2016b) is chosen to determine SMR based on calculation of the lower quantile (p = 0.25) of all slopes with *r^2^* > 0.9 (the threshold level is reduced because of the high rate of background respiration, coupled with a low ratio between mass of male guppies and the volume of chambers, both of which seriously reduced the linearity of corrected data). Note, if any of ‘calcSMR’ methods are applied, then slopes cannot be plotted individually in the module ‘QC of slopes’. Finally, we insert information about the body density of fish and click the button ‘Calculate and plot’. Three graphs are plotted automatically in the graphical module: percentage rate of background respiration, absolute and mass-specific SMR. The last step is to save results as a .csv or .txt file (a log file with all commands applied during the analysis is also logged automatically in the same folder).

Results of the analysis show that background respiration represents 57.1-89% of the total oxygen consumption (Fig. 7A), despite daily cleaning of chambers and sensors, weekly changing 50% of water volume in the respirometry system and constant usage of the UV filter. Absolute SMR varies depending on mass of fish, while the mean mass-specific SMR is 185.5 mgO_2_/kg/h. Thus, mean SMR is overestimated by approximately 2.8 times for females and 6.7 times for males (Fig. 7B), when not corrected for background respiration.

**Figure 7.**
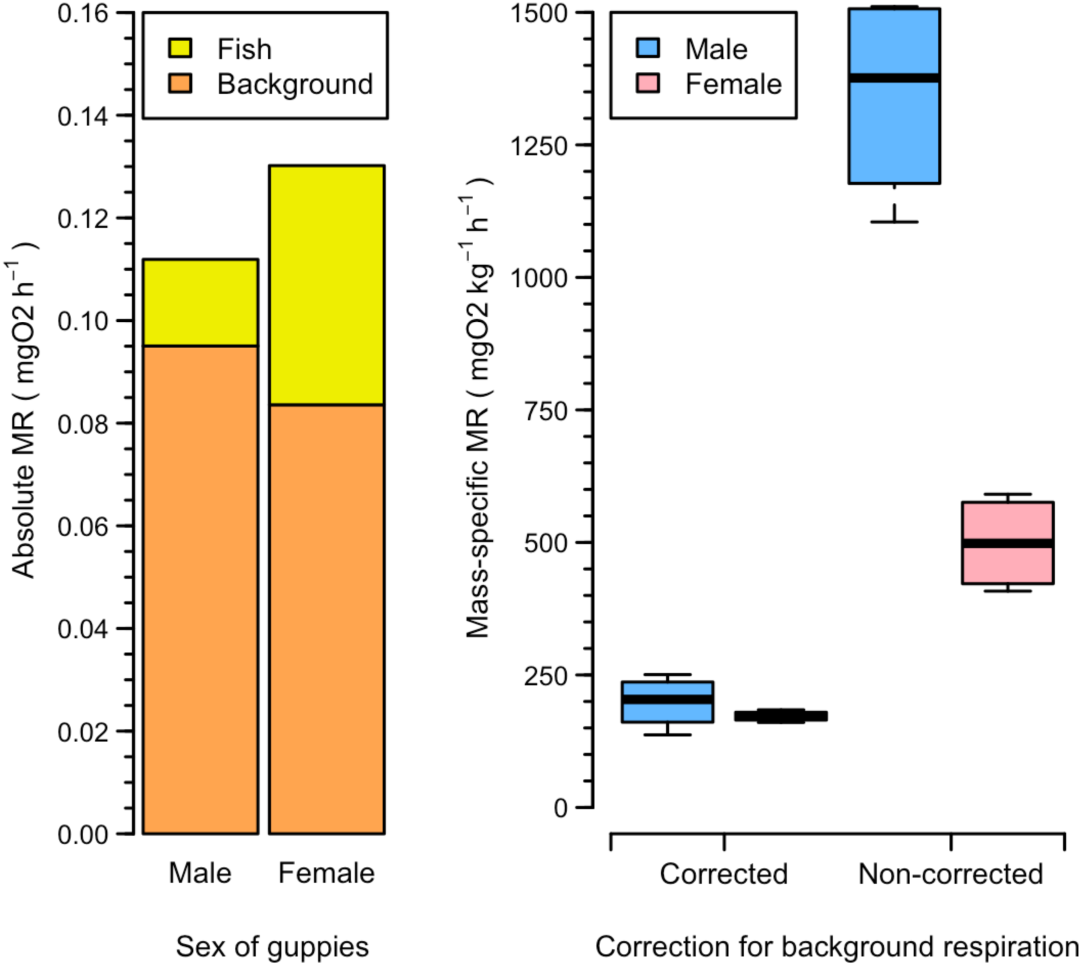
(A) Absolute SMR of guppies versus background respiration. (B) Difference in corrected and
non-corrected mass-specific SMR.

## 4 Conclusions

Our case studies demonstrate that neglecting mechanical problems and microbial growth in a respirometry system can lead to serious underestimation and/or overestimation of metabolic rate. FishResp software, used together with a proper experimental design (Clark et al., 2013; Svendsen et al., 2016b), represents a simple and efficient approach to control both chamber and background respiration effects in the processing of raw respirometry data. To make FishResp a universal tool, we have developed both the R package (available on CRAN), and its cross-platform and user-friendly graphical implementation under open source licenses. Both can import raw data from the majority of respirometry systems available on the market. FishResp can be adjusted for different experimental designs: the software allows users to choose among various options for detection of mechanical problems, correction of background respiration, and calculation of metabolic rate. Therefore, FishResp provides a universal and straightforward tool that can improve the quality of metabolic rate estimation in fishes and other aquatic animals. Last but not least, the ‘Extensions module’ in the GUI version of FishResp represents a platform for graphical implementation of other R packages, scripts and functions available in open-access/published. Thus, FishResp has potential to become a community-driven software in the field of aquatic respirometry.

## 5 Acknowledgements

We thank Artur Zaripov and Roman Kovalev for significant contribution to GUI development, Federico Calboli and Nicholas Carey for helpful advices and testing the R package, Niclas Kolm and Alexander Kotrschal for providing the guppy data. Financial support from Oskar Öflund Foundation (SM), Alfred Kordelin Foundation (SM), Helsinki Institute of Life Sciences (#797011029 to JM) and Academy of Finland (# 259944 to RJSM; # 292737 to JM) is gratefully acknowledged.

## 6 Data Availability

The R package and data are available on the CRAN repository (http://cran.r-project.org/package=FishResp). R scripts for running both case studies are provided in supplementary files S1 & S2. Application of functions which have not been reviewed in the case studies are shown in supplementary file S3. The official website (https://fishresp.org) will host all information related to FishResp, including user manuals, tutorial videos, FAQ, a forum and a blog with the latest updates of the software.

## 7 Authors’ Contributions

SM and RJSM conceived the ideas and designed the original R functions. SM developed the R package and software; All authors led the writing of the manuscript and contributed critically to all drafts, and gave final approval for publication.

